# Genomic Informational Field Theory (GIFT) to identify genetic associations of a complex trait using a small sample size

**DOI:** 10.1101/2025.08.15.670531

**Authors:** Panagiota Kyratzi, Scott Gadsby, Edd Knowles, Patricia Harris, Nicola Menzies-Gow, Jonathan Elliott, Andras Paldi, Jonathan Wattis, Cyril Rauch

**Author notes:** Correspondence: Cyril Rauch.

## Abstract

Genome-wide association studies (GWAS) are commonly used to investigate the genetic basis of complex traits. However, to be adequately powered they typically require large sample sizes to provide precise inferences. To address this challenge, this paper introduces GIFT, a novel data analytic method that enhances the power of genetic analyses, enabling the use of smaller datasets without compromising precision. In a small cohort of 157 ponies, GIFT was applied to examine the complex trait of “height at withers”, comparing its performance to traditional GWAS. GIFT enabled the identification genetic loci linked to insulin physiology validating, in turn, a long-standing hypothesis that “height at withers” is associated with insulin physiology in equids, potentially promoting equine metabolic syndrome (EMS). By redefining correlations between single nucleotide polymorphisms (SNPs), GIFT provides new insights into linkage disequilibrium and reveals underlying gene network structures. This, in turn, enables the distinction between core and peripheral genes within these networks. By reducing the time and cost associated with large-scale genotype–phenotype mapping studies without sacrificing statistical robustness, GIFT broadens access to quantitative genetic research, allowing smaller-scale studies to investigate the genetic architecture of complex traits with greater resolution.

**NEW & NOTEWORTHY:** Inferring genotype–phenotype associations typically require large sample sizes, limiting many genetic studies. GIFT, a novel data analytics tool, overcomes this by enabling accurate association mapping in small datasets. We further show how GIFT’s extended framework infers linkage disequilibrium and gene networks, distinguishing core from peripheral genes involved in complex traits. This advancement enhances understanding of biological architecture and enables high-resolution genetic research in limited cohorts, offering a powerful, cost-effective alternative to traditional large-scale approaches.

## INTRODUCTION

Identifying associations between phenotype and genotype is the fundamental basis of genetic analysis and GWAS is currently the method of choice for mapping genotype to phenotype in large populations. It is expected that large-scale GWASs will elucidate the genetic basis of complex traits. The need for “large-scale” GWAS arises from the statistical requirements of the method requiring very large populations to produce valid inferences when effect sizes are small. A prime example of large-scale GWAS applied to a complex trait is the genetic analysis of the phenotype “standing height” in humans. A recent GWAS meta-analysis involving an unprecedented sample size of 5.4 million individuals successfully produced a saturated map of genetic variants associated with human height (1). Although this represents a landmark achievement in genetics, it also highlights the limitations of GWAS, whose success relies heavily on data acquisition to generate adequately powered studies, which can be a limitation to individual investigations or non-consortium science. While human genetic studies benefit from substantial funding and large-scale GWAS capabilities, research on non-human species, especially those facing extinction, often lacks the necessary resources to conduct similar studies (2). As a result, there is a need to explore alternative methods, increasing the investigative power of biological datasets when sample sizes are small.

GWAS is based on the statistical framework developed by Fisher more than 100 years ago. It partitions genotypic values and performs linear regression of phenotype on marker allelic dosage. Regression coefficients estimate the average allele effect sizes, and the regression variance is the additive genetic variance due to the locus. However, there is ongoing debates about whether the statistical method underpinning GWAS is optimal for understanding complex traits (3, 4).

In GWAS, statistical associations rely on frequency or count plots like bar charts or histograms, which categorize data into bins. These plots, known as distribution density functions (DDFs), form the basis of frequentist probability, determining key statistical measures such as averages and variances that are used to infer genotype-phenotype associations. However, this approach involves the loss of significant amounts of information from datasets. To comprehend this point, consider a population where phenotypic data are measured with high precision, making each data point unique. When grouped into bins to form DDFs, individual distinctions are lost since it is not possible to differentiate data within a given category, even though the data were initially differentiated. While averages and variances derived from DDFs appear informative, they are inherently based on this reduced representation limiting access to fine-grained information. Increasing the sample size might improve the precision of statistical summaries, i.e., average and variance, but it does not recover the lost individual details.

This raises a fundamental question: Is more data always beneficial if the method inherently disregards fine-grained information?

GIFT is a novel data analytics method designed to overcome this conceptual limitation by analyzing the full biodiversity of data without requiring data categorization. Through theoretical simulations (5, 6) and analysis of various datasets with sample sizes comparable to those used in GWASs (7), GIFT has been shown to enhance both investigative and discriminative powers of genetic studies. However, the ability of GIFT to operate effectively with smaller sample sizes and provide essential genetic information has not yet been shown.

Herein, using a small cohort of 157 ponies and focusing on the complex trait “height at withers” the investigative powers of GWAS and GIFT are compared. Several reasons motivated the focus on “height at withers”.

First, like “standing height” in humans, “height at withers” in equids is a classic complex trait positively selected during domestication and correlated with other morphometric traits (8–10). With a well-established heritability, GWASs have investigated its genetic background (9, 11–24). Thus, GIFT-derived variants can be benchmarked against GWAS results in the equine species and compared to similar phenotype, i.e., “body height”, across different species.

Second, “height at withers” is suspected to be associated with equine metabolic syndrome (EMS) (11). EMS describes a cluster of metabolic disturbances predisposing horses to laminitis, a condition where insulin dysregulation plays a central role (11, 16, 25, 26). While a GWAS identified HMGA2 (High mobility group AT-hook protein 2) and IRAK3 (Interleukin-1 receptor-associated kinase 3) as associated with “height at withers” in a cohort of 236 ponies, no other genes were linked to insulin physiology (11). However, since a retrospective correlation was identified between blood insulin concentrations and HMGA2 allelic dosages (11), it was hypothesized that GIFT could reveal further genetic insights into the relationship between insulin physiology and “height at withers” (27).

Third, it is postulated that complex traits can be omnigenic involving core and peripheral genes (28). Validating this new paradigm using GWAS would necessitate large sample sizes to detail any information on gene networks. However, since GIFT provides a more detailed genetic analysis, it was hypothesized that redefining SNP and gene pairwise correlations using GIFT could offer a fresh perspective on linkage disequilibrium (LD) and gene networks informing, in turn, on the potential validity of the omnigenic paradigm.

The results herein demonstrate that GIFT’s approach is adequate when small sample sizes are used, yielding new insights into the genetic architecture of complex traits.

## MATERIALS AND METHODS

### Data collection

Data from a study performed by Knowles *et al*. (conducted with ethical approval from the Royal Veterinary College Animal Welfare and Ethical Review Board URN 2015-5128) were re-used in which blood collection was performed and the “height at withers” measured in a number of different breeds (25). In order to remain coherent with the study by Norton *et al* (11), Shetland ponies were removed from the current study resulting in a final sample size of 157 ponies

### Equine genome sequencing

Blood samples collected (buffy coats) were used for DNA extractions (LGC Biosciences) and genotyping individuals using the Axiom™ Equine Genotyping Array (MNEC670, Cambridge Genomic Services). Further to Axiom Analysis Suite following best practices pathway, a total of 478498 quality controlled single nucleotide polymorphisms were identified.

### Phenotype ‘height at withers’

Measurements of “height at withers” obtained in (25) were pre-analysed. A Kolmogorov-Smirnov test performed against raw data for the height returning a p-value of 0.05101 suggested that raw data are not normally distributed, requiring the need to pre-correct data for fixed effects. Phenotypic data were then pre-corrected for the fixed effects of breed and sex, as well as for relatedness between individuals using the kinship matrix (K) extracted using GEMMA. Using R-studio (GMMAT package) the following linear model was used, 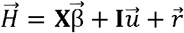, where 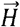 is the vector of individual height, **X** is the designed matrix for fixed effects, 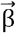 is the vector of coefficients for fixed effects, **I** is the identity matrix, 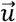 is a vector of normally distributed random effect linked to the kinship matrix **K** defined by, 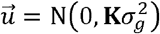,where 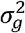 is the genetic variance and 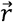 is the residual vector. With this correction, a Kolmogorov-Smirnov test was performed against the residual values for the height providing an acceptable p-value of 0.836 demonstrating, in turn, that the components of 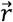 are quasi-normally distributed and usable by GWAS.

### GWAS

The genotype dataset was filtered using PLINK (HWE p-value threshold of 10^-6^, call rate for genotypes of 10% and a MAF of 10%), the number of independent SNPs (N_lnd-SNPs_ = 44412) was determined using BCFTOOLS (r^2^-threshold=0.1) and the GWAS Manhattan plots, linked to the determination of the GWAS p-values (p_GWAS_), were obtained using GEMMA. The threshold of significance for GWAS was determined using 99% and 95% Bonferroni corrections, numerically given by, −Log(0.01/N_Jnd-SNPs_) =6.647 and −Log(0.05/N_Jnd-SNPs_) = 5.9485, respectively.

### GIFT

The same genotype and phenotypic residuals, as filtered by GWAS, were used for GIFT. Regarding the representation of GIFT, upon selecting a SNP the different genotypes were assigned arbitrary values of +1, 0, and −1, respectively. More specifically, the assignment of values +1, 0, and −1 were done as a function of the base pairs as follows: AA=TT=+1, GG=CC=−1, and 0 otherwise. The genetic microstates +1, −1 and 0 are respectively color coded in red, blue and grey in the figures to facilitate the representation of genetic paths. With this convention, any barcode can be represented by a string of numbers from which a GIFT analysis can be performed providing a level of significance noted p_GJFT_ for the SNP considered, as explained in (7) and further developed mathematically in (27). The level of significance (p_GJFT_) can then be plotted using a Manhattan plot and the same significance thresholds determined using 99% and 95% Bonferroni correction as in GWAS. For readers unfamiliar with GIFT, Supplemental_Text_S1.doc introduces the methodology underpinning GIFT.

### Enrichment and correlation analyses

Enrichment analyses were performed using STRING, Uniprot, Ensembl, Gephi and the GWAS catalog (29– 33) as cited in the text. Pearson correlations between genetic paths were performed using R-studio and the measure of centrality using Matlab.

## RESULTS

### Manhattan Plots using GWAS or GIFT

GIFT is a method for inferring genotype–phenotype associations without relying on data categorization or grouping, challenging the common perception arising from DDF-based statistical approaches in genetics that robust and precise inferences are inherently dependent on large sample sizes.

Measurements of “height at withers” obtained by Knowles et al. (25) were pre-corrected for fixed effects and relatedness. Further to this correction, phenotypic residual values were demonstrated to be adequately normally distributed using Kolmogorov-Smirnov test and usable by GWAS and GIFT (Supplemental_Figure_S2.doc). Association analyses using GWAS and GIFT based on the same phenotypic residual values were then performed and plotted under the form of Manhattan plots. Fig.1 presents the Manhattan plots for both GWAS (Fig.1A) and GIFT (Fig.1B). Concentrating on the highest threshold levels for GWAS and GIFT, the significant SNPs were then mapped to the reference horse genome assembly (EquCab3.0) available on Ensembl to retrieve the corresponding gene Id and gene name when available. The results summarized in Table 1 and Table 2 clearly demonstrate that SNPs located on chromosome 6 are highly significant. However, while GWAS identifies 5 significant SNPs associated with the HMGA2 gene, GIFT detects up to 18 significant SNPs for the same gene on chromosome 6. Notably, GIFT also highlights significant SNPs on chromosome 2 that are not detected by GWAS. Figures 1C and 1D provide a comparative visualization of two representative SNPs using both GWAS and GIFT. Figure 1C shows a significant SNP on chromosome 6 identified by both methods, whereas Figure 1D presents one significant SNP on chromosome 2, uniquely identified by GIFT. Interestingly, the SNP on chromosome 2 exhibiting a sigmoidal genetic path (Figure 1D, right panels) lacks a traditionally defined gene effect size (Figure 1D, left panel).

**Figure 1:**
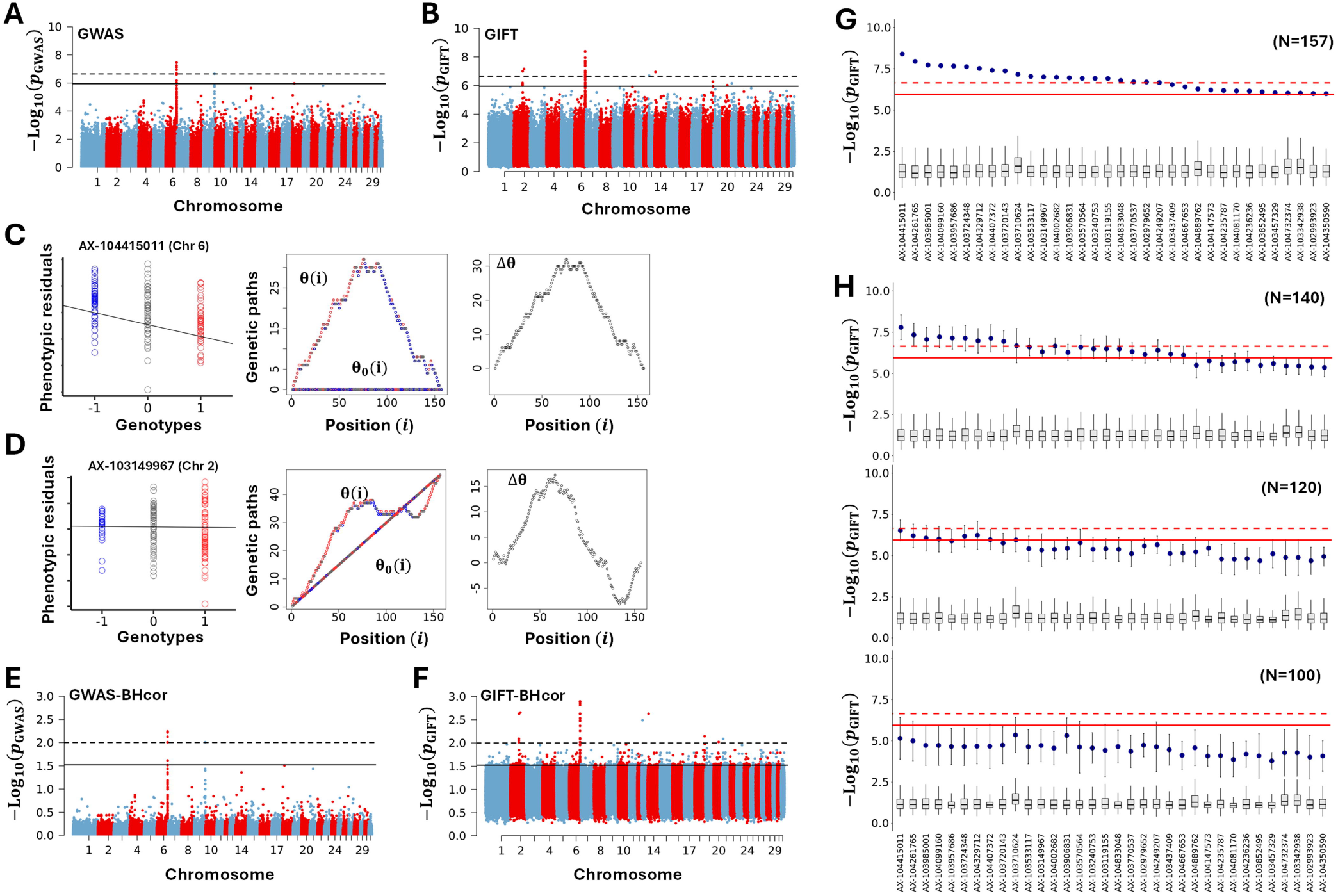
Comparison of GWAS and GIFT results for “height at withers”. **(A)** GWAS Manhattan plot. The vertical dashed and solid lines represent the 99% and 95% Bonferroni correction thresholds, respectively.**(B)** GIFT Manhattan plot. The vertical dashed and solid lines represent the 99% and 95% Bonferroni correction thresholds, respectively. **(C)** GWAS (left panel) and GIFT (middle and right panels) representations of SNP-Id: AX-104415011 (Chr6, position: 82439707) significant for both GWAS and GIFT with respective significances, −log(p_GWAS_) = 7.246 and −log(p_GJFT_) = 8.39. The GIFT representation displays a parabolic genetic path that is characteristic of GWAS-derived associations. θ(*i*) represents the cumulative sum of genotypes (SNP-Id AX-104415011) ordered according to the magnitude of the trait “height at withers” after correction for fixed effects and relatedness. Importantly, θ(*i*) is defined based on phenotypic information. In this context the variable “*i*” denotes the position in the ordered list. θ_0_(*i*) is obtained by calculating the average genetic path derived from an infinite number of random permutations of genotypes. Because random permutation removes any relationship with the phenotype, θ_0_(*i*) represents the null expectation (i.e., absence of association) within the GIFT framework. The difference between these two quantities, Δθ(*i*), is then used to derive significance levels for genotype–phenotype associations (for more details see Supplementary_Text_S1 or (7, 27)). **(D)** GWAS (left panel) and GIFT (middle and right panels) representations of SNP-Id: AX-103149967 (Chr2, position: 82564015) only found significant by GIFT, −log(p_GJFT_) = 6.999. The GIFT representation displays a sigmoidal genetic path that cannot be detected through GWAS-derived associations. **(E)** Manhattan plots for GWAS after correcting for false-discovery rates using the Benjamini-Hochberg procedure. **(F)** Manhattan plots for GIFT after correcting for false-discovery rates using the Benjamini-Hochberg procedure. **(G)** To demonstrate the absence of spurious association with GIFT, the microstates of each significant SNP determined by GIFT were permutated randomly 1000 times and the statistics of their −*log*_l0_(*p*_*GJFT*_) values displayed using a box plot and compared to their initial significance level (blue disk). The red dashed and continuous lines correspond, respectively, to 99% and 95% Bonferroni correction thresholds. **(H)** To determine GIFT’s robustness, the original dataset (N=157) was randomly subsampled to N=140, 120, and 100. This procedure was repeated 10 times for each subsample to compute the mean and standard deviation of −*Log*_l0_(p_GJFT_) value for the most significant SNPs (blue disk). The ordering of SNPs on the x-axis is the same as in (E). Additionally, the microstates were permutated randomly 1000 times and the statistics of their −*log*_l0_(*p*_*GJFT*_) values displayed using a box plot.Supplemental_Data_S3.xls provides the correspondence between the Axiom− SNP annotations, typically ‘AX-…’, and the chromosomal positions of the significant SNPs for GWAS and GIFT. (Chr: chromosome).

**Table 1.** List of significant genes detected by GWAS ordered by p-values.

| **GWAS (99% Bonferroni correction)** |  |  |  |  |  |  |
| --- | --- | --- | --- | --- | --- | --- |
| **Chr.** | **Gene-Id** | **Gene name** | **Biotype** | **#SNPs detected** | **$\langle -\log(p_{\text{GWAS}}) \rangle$** | **$\sigma_{-\log(p_{\text{GWAS}})}$** |
| 6 | ENSECAG00000040252 | HMGA2 | protein | 5 | 7.129 | 0.298 |
| **GWAS (95% Bonferroni correction)** |  |  |  |  |  |  |
| **Chr.** | **Gene-Id** | **Gene name** | **Biotype** | **#SNPs detected** | **$\langle -\log(p_{\text{GWAS}}) \rangle$** | **$\sigma_{-\log(p_{\text{GWAS}})}$** |
| 6 | ENSECAG00000040252 | HMGA2 | protein | 5 | 7.129 | 0.330 |
| 6 | ENSECAG00000057296 | NA | lncRNA | 2 | 6.317 | 0.418 |
| 6 | ENSECAG00000051733 | NA | lncRNA | 1 | 6.021 | NA |
| 18 | ENSECAG00000017122 | NCKAP5 | protein | 1 | 5.973 | NA |

When multiple significant SNPs were detected on a gene, $\langle -\log(p_{\text{GWAS}}) \rangle$ represents the average p-value calculated and, $\sigma_{-\log(p_{\text{GWAS}})}$ represents the standard deviation of p-values calculated. Supplemental\_Data\_S3.xls provides detailed information on the SNPs involved. (Chr.: Chromosome).

**Table 2.** List of significant genes detected by GIFT ordered by p-values.

| **GIFT (99% Bonferroni correction)** |  |  |  |  |  |  |
| --- | --- | --- | --- | --- | --- | --- |
| **Chr.** | **Gene-Id** | **Gene name** | **Biotyp e** | **#SNPs detected** | **$\langle -\log(p_{\text{GIFT}}) \rangle$** | **$\sigma_{-\log(p_{\text{GIFT}})}$** |
| 6 | ENSECAG0000005729<br>6 | NA | lncRNA | 2 | 7.783 | 0.230 |
| 6 | ENSECAG0000005173<br>3 | NA | lncRNA | 1 | 7.620 | NA |
| 6 | ENSECAG0000004025<br>2 | HMGA2 | protein | 14 | 7.161 | 0.383 |
| 2 | ENSECAG0000002208<br>9 | CBR4 | protein | 1 | 6.999 | NA |
| 2 | ENSECAG0000002312<br>2 | PALLD | protein | 1 | 6.999 | NA |
| **GIFT (95% Bonferroni correction)** |  |  |  |  |  |  |
| **Chr.** | **Gene-Id** | **Gene name** | **Biotyp e** | **#SNPs detected** | **$\langle -\log(p_{\text{GIFT}}) \rangle$** | **$\sigma_{-\log(p_{\text{GIFT}})}$** |
| 6 | ENSECAG0000005729<br>6 | NA | lncRNA | 2 | 7.783 | 0.230 |
| 6 | ENSECAG0000005173<br>3 | NA | lncRNA | 1 | 7.620 | NA |
| 2 | ENSECAG0000002208<br>9 | CBR4 | protein | 1 | 6.999 | NA |
| 2 | ENSECAG0000002312<br>2 | PALLD | protein | 1 | 6.999 | NA |
| 6 | ENSECAG0000004025<br>2 | HMGA2 | protein | 18 | 6.966 | 0.512 |
| 18 | ENSECAG0000002432<br>5 | KIAA201<br>2 | protein | 1 | 6.270 | NA |
| 18 | ENSECAG0000002469<br>3 | SUMO1 | protein | 1 | 6.270 | NA |
| 2 | ENSECAG0000005042<br>8 | NA | lncRNA | 3 | 6.093 | 0.09 |
| 20 | ENSECAG0000001839<br>3 | BTBD9 | protein | 2 | 6.022 | <0.001 |
| 6 | ENSECAG0000000672<br>1 | TMBIM4 | protein | 1 | 5.989 | NA |

When multiple significant SNPs were detected on a gene, $\langle -\log(p_{\text{GIFT}}) \rangle$ represents the average p-value calculated for the gene and, $\sigma_{-\log(p_{\text{GIFT}})}$ represents the standard deviation of p-values calculated. Supplemental\_Data\_S3.xls provides detailed information on the SNPs involved. (Chr.: Chromosome).

While the Bonferroni correction provides strong protection against type I errors, it is highly conservative, particularly in GWAS using relatively small sample sizes, and may therefore miss biologically relevant associations. In contrast, the Benjamini–Hochberg procedure, which controls the false discovery rate (FDR) using the expected proportion of false positives among significant results, is preferred for small genetic population studies and exploratory analyses, as it maximizes discovery while controlling error rates.

Accordingly, p-values from both GWAS and GIFT were adjusted using the Benjamini–Hochberg procedure and compared. As shown in Fig. 1E and Fig. 1F for GWAS and GIFT, respectively, a large proportion of the signals detected by GIFT remain significant after FDR correction (adjusted p-value < 0.05). This result indicates that, under the same error-control criteria, GIFT is more sensitive than GWAS in identifying significant associations in studies with small sample sizes.

To further confirm that the significant SNPs identified by GIFT in Fig.1B result from the unique organization of their microstates rather than random chance, the microstates of each significant SNP from Fig.1B were randomly permuted 1,000 times. After each permutation, the significance level, i.e., the value of −Log_l0_(p_GJFT_), was determined. The statistical summary of these significant values obtained from 1,000 random permutations was presented as a boxplot (Fig.1G). These random permutations represent the distribution expected under a null hypothesis in which no genotype–phenotype association exists for the SNP considered. These values were then compared to the original significance value of the same SNP and to the Bonferroni corrections applied, as determined in Fig.1B, and represented as a blue dot in Fig.1G. Fig.1G confirms that GIFT’s identification of genotype–phenotype associations is not due to random chance.

Finally, to assess whether GIFT can operate effectively with smaller sample sizes, the original dataset (N=157) was randomly subsampled to N=140, 120, and 100. For each subsample, the values of −Log_l0_(p_GJFT_) corresponding to the most significant SNPs in Fig.1B were recalculated. This procedure was repeated 10 times for each new sample size to compute the average and standard deviation of −Log_l0_(p_GJFT_) values obtained for the top SNPs (Fig.1E). Additionally, for each subsampling the microstates were randomly permuted 1,000 times to extract an average value for −Log_l0_(p_GJFT_). As each subsampling was performed 10 times the statistics (average and standard deviation) of −Log_l0_(p_GJFT_) values upon subsampling and random permutation of microstates were reported as a boxplot for each SNP. As shown in Fig.1F, although a decrease in the magnitude of −Log_l0_(p_GJFT_) is clearly observed with smaller sample sizes, there is no overlap of standard deviations obtained between the subsampling and the null hypothesis.

### Biological relevance of genes determined by GWAS and GIFT

Focusing on the 99% Bonferroni correction threshold, the GWAS identified HMGA2, a gene associated with the trait “body size” in several animal species, including ponies, sheep, dogs, and cattle (11, 34–36). When the Bonferroni threshold was relaxed to 95%, an additional gene, NCKAP5 (NCK-associated protein 5), was identified. Interestingly, although NCKAP5 does not appear among the significant genes listed by GIFT above the 5% Bonferroni correction threshold, its significance value, −Log(p_GJFT_), remains high, at 5.301. With regard to GIFT and using the lowest threshold, the significant genes detected were HMGA2, CBR4 (carbonyl reductase 4), PALLD (palladin), KIAA2012, SUMO1 (small ubiquitin-related modifier 1), BTBD9 (BTB domain–containing 9), and TMBIM4 (transmembrane BAX inhibitor motif–containing protein 4).

Given the limited number of extensive genetic studies in equids compared to humans, the relevance of the gene names listed in Tables 1 and 2 was initially assessed using “body height” as a search term in the GWAS Catalog (29). All genes listed in Tables 1 and 2, with the exception of SUMO1, have previously been significantly associated with “body height” in humans (1).

In a second step, the genes listed in Tables 1 and 2 were cross-referenced in PubMed to determine whether their reported functions are related to physiological processes associated with insulin action, insulin secretion, glucose uptake and homeostasis, energy metabolism, and metabolic disorders such as diabetes. Given the limited number of molecular biology studies in equids, this investigation was extended to other species.

HMGA2 (High Mobility Group AT-hook 2) is a non-histone chromatin-associated protein that binds AT-rich DNA sequences to modulate gene expression, acting as an architectural transcription factor (37), including the expression of peroxisome proliferator-activated receptor γ (PPARγ) leading to the expression and release from adipose tissue of adiponectin, an insulin-sensitizing adipokine (38). While HMGA2 itself has not been directly linked to metabolic dysregulation, in obesity and type 2 diabetes the impairment of PPARγ activity selectively downregulates insulin-sensitizing genes, most notably adiponectin, thereby contributing directly to systemic insulin resistance (39).

A multi-omic mapping of human pancreatic islet stress responses has identified NCKAP5 as a gene dynamically regulated by both endoplasmic reticulum (ER) stress and pro-inflammatory cytokines, two pathophysiological stressors implicated in type 2 diabetes (T2D) pathogenesis (40).

CBR4, a member of the carbonyl reductase family, regulates fatty acid biosynthesis through its interaction with fatty acid synthase (FASN), a multifunctional enzyme central to energy storage and signal transduction. FASN has been implicated in activation of the PI3K/AKT/mTOR signaling axis, which governs cellular growth and metabolism (41). Mechanistically, CBR4 engages the ubiquitin-proteasome pathway to promote the degradation of FASN, resulting in reduced FASN protein levels and subsequently diminished AKT (protein kinase B) activation (42, 43). This cascade of events ultimately leads to the inhibition of the mTOR (mechanistic target of rapamycin) signaling pathway. Given that mTOR is directly activated by insulin through the canonical PI3K/AKT cascade, this regulatory axis positions CBR4 as a modulator of insulin-mediated signals that integrate cellular energy status (43).

PALLD (Palladin), a cytoskeletal protein, is essential for active actin remodelling and the formation of membrane ruffles (44), a process integral to insulin signalling due to its role in the membrane translocation of specific GLUT family members (45). Furthermore, a genome-wide functional screen identified PALLD as a novel negative regulator of insulin signaling, demonstrating that its overexpression inhibits insulin-stimulated AKT phosphorylation and reduces insulin receptor (IR) abundance, while its knockdown produces the opposite effects (46).

SUMO1 is a central component of the SUMOylation pathway. This post-translational modification process critically regulates metabolic homeostasis by modifying key transcription factors that influence lipid biosynthesis, adipogenesis, and energy balance. SUMO1 covalently attaches to specific lysine residues on target proteins, thereby modulating their stability, subcellular localisation, and transcriptional activity in response to metabolic cues (47, 48). A pivotal target of SUMO1 is the nuclear receptor PPARγ, a master regulator of insulin sensitivity (49). Functional studies using SUMO1-knockout mice revealed that these animals gain less weight on a high-fat diet and show attenuated expression of PPARγ target genes, demonstrating that SUMO1 is required for full PPARγ transcriptional activity (50). Collectively, these findings establish SUMO1 as a critical modulator of transcriptional networks governing adipogenesis, lipid metabolism, and energy balance, with direct implications for insulin sensitivity and the pathogenesis of metabolic disorders.

Mechanistic studies have identified BTBD9 as a novel component of the insulin/insulin-like growth factor (IGF) signaling pathway, where it functions as a negative regulator. Specifically, BTBD9 overexpression significantly elevates levels of the forkhead box O (FOXO) transcription factor while decreasing AKT levels, establishing a mechanistic link to the canonical insulin/IGF signaling cascade (51, 52). Interestingly, human subjects with elevated blood manganese levels exhibited decreased BTBD9 mRNA expression, suggesting environmental regulation of this modulator (52). These findings position BTBD9 as a key regulator of IGF signaling connecting BTBD9 to fundamental growth factor signaling pathways involved in body growth and development.

TMBIM4 was identified as a novel candidate gene for type 2 diabetes (T2D) through a multi-tissue gene expression analysis that compared transcriptomic data from T2D patients and controls across 21 experimental groups sourced from the Gene Expression Omnibus (GEO) database. The gene was prioritized based on two complementary criteria: it exhibited a high percentage of differential expression between diabetic and non-diabetic individuals across multiple tissues, and it was located within ±1 Mb of a known T2D susceptibility SNP (rs1531343), suggesting it may be the functional effector gene through which this genetic variant exerts its pathogenic effect. Expression analysis revealed that TMBIM4 was predominantly upregulated in skeletal muscle but downregulated in liver tissue of T2D patients, indicating tissue-specific dysregulation in key insulin-responsive organs (53).

### GIFT’s determination of LD

Altogether, GIFT’s results strongly suggest that gene associated with the phenotype “height at withers” are also related to metabolic dysregulations.

Notably, the genes listed in Table 2 are located on different chromosomes, indicating that GIFT’s pairwise SNP correlations may offer novel insights into linkage disequilibrium (LD) possibly highlighting how genes may interact together.

At the core of GIFT lies the concept of the genetic path, represented by Δθ in Fig.1C and Fig.1D (right panels) for each SNP in the genome. The overall pattern of Δθs reflects the ordering of genetic microstates when the sample is ordered by phenotypic residual values. Consequently, SNPs similarly related to the phenotype exhibit comparable large Δθ patterns. Conversely, SNPs that are unrelated to the phenotype give small and noisy Δθ curves (see supplemental_Text_S1). This property is essential to determine precise correlations between loci, which can be compared to that of usual LD definition. An example will illustrate this point.

Using GIFT, HMGA2 contains 18 significant SNPs (Table 2) and it is therefore possible to determine the local LD in the portion of genome that defines this protein. The measure of LD commonly used, a.k.a. the r^2^ measure of LD, is given by (54):

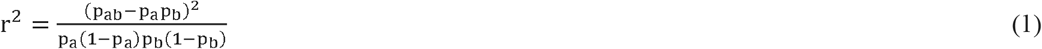

where p_ab_ is the frequency of haplotypes having allele ‘a’ at locus 1 and allele ‘b’ at locus 2. As the square of a correlation, r^2^ can range from 0 to 1 as p_a_, p_b_ and p_ab_ vary. In Fig.2A, the genetic paths θ(i) and θ_0_(i) pertaining to the 18 SNPs belonging to HMGA2 (Table 2), enumerated from 1 to 18 in the figures, are represented. In Fig.2B the r^2^ measure of LD for HMGA2 is determined where the enumeration of SNPs (from 1 to 18) is given next to their SNP-Id on the diagonal. Fig.2B demonstrates that while the r^2^ measure of LD for HMGA2 is very patchy, strong correlations exist between some SNPs. It is interesting to note from Fig.2B that while the correlation between SNPs 1 and 5, marked by a white star in Fig.2B is relatively strong, the paths are anti-symmetrical. This anti-symmetry arises because, for SNP 1, the path begins with ‘−1’ (blue) microstates and ends with ‘+1’ (red) microstates, whereas the opposite is observed for SNP 5. This antisymmetric effect is not reflected in equation (1), as it considers the entire population to compute bulk probabilities (p_a_, p_b_, p_ab_), meaning that the notion of ordering does not influence the result from equation (1).

**Figure 2:**
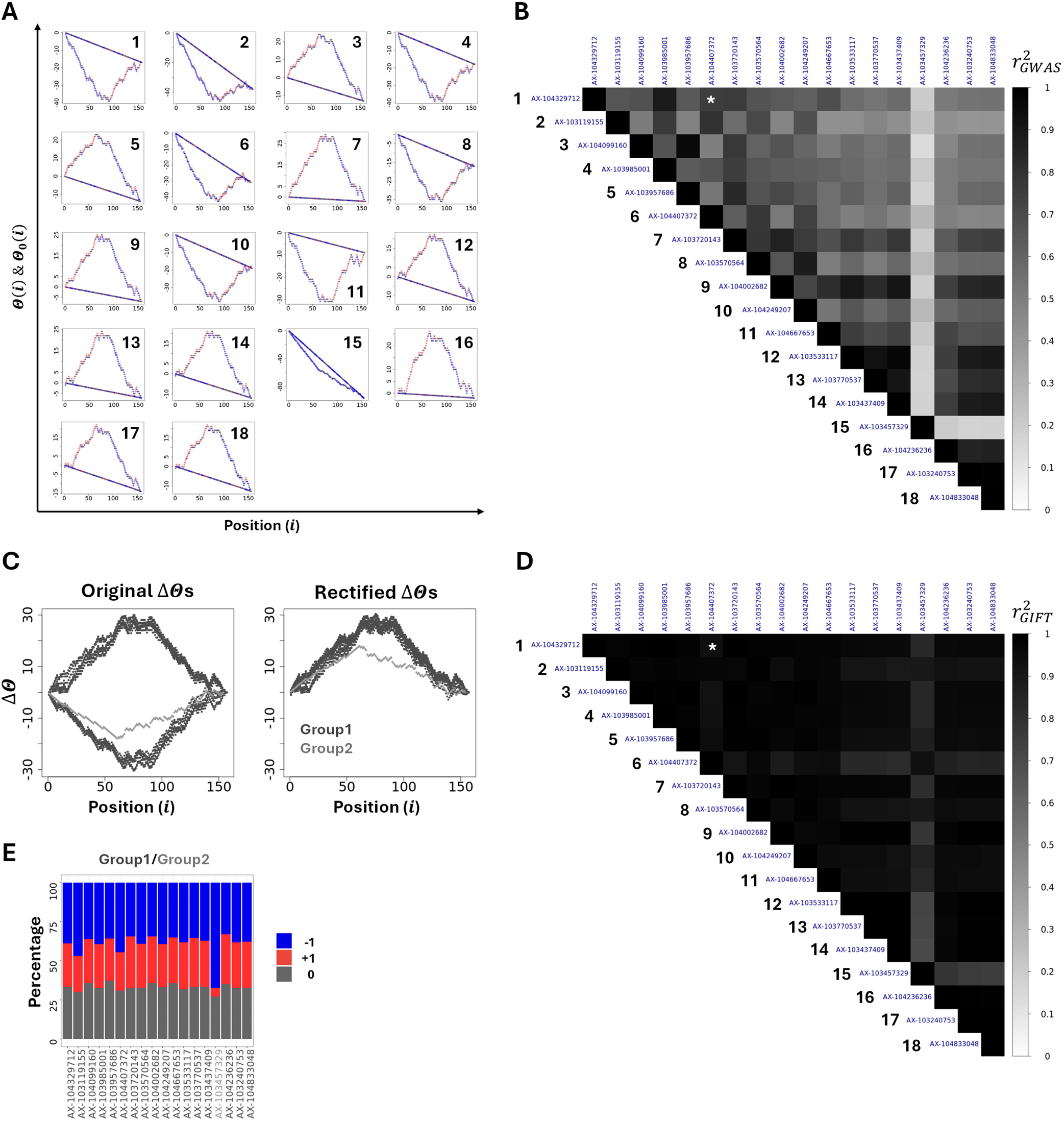
Genetic Paths and Linkage Disequilibrium (LD) for HMGA2. **(A)** Plot of genetic paths, θ(i) and θ_0_(i), for the first 18 SNPs associated with HMGA2, numbered from 1 to 18. **(B)** r^2^ measure of linkage disequilibrium (LD) for HMGA2. The first 18 SNPs (as shown in (A)) are enumerated from 1 to 18, with their SNP-Ids displayed along the diagonal. **(C)** Genetic trajectories (Δθs) of all SNPs within HMGA2 are plotted (left panel) and rectified (right panel), revealing the presence of two SNP types (coloured in light and dark grey). **(D)** 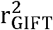 measure quantifies pairwise correlations between SNPs within HMGA2. **(E)** The SNPs identified exhibit strong symmetries when analysed in terms of microstate composition. Supplemental_Data_S4.xls provides the correlation values computed.

This highlights that the initial and arbitrary assignment of microstates (‘+1’ and ‘−1’) is not fundamental for the r^2^ measure of LD.

To develop a novel measure of LD based on genetic paths, while maintaining the core principles of equation (1), the genetic paths in Fig.2A can be rectified and overlapped to reveal their underlying patterns. This is possible by swapping the initial assignment of microstates (‘+1’ and ‘−1’). Note that as the level of significance calculated by GIFT relies on the amplitude of Δθ (see supplemental_Text_S1), changing the initial assignment of microstates, e.g., swapping the microstates ‘+1’ and ‘−1’, does not change the level of significance calculated. In the left panel of Fig.2C, the genetic paths (Δθs) of all SNPs within HMGA2 are plotted. In the right panel of Fig.2C, these paths are rectified and overlapped, revealing the presence of two distinct types of genetic paths (light grey and dark grey). Since the key to inferring associations with the phenotype lies in the overall pattern of these genetic paths, a refined measure of LD as described by GIFT can be formulated simply using Pearson’s correlation formula in the following form:

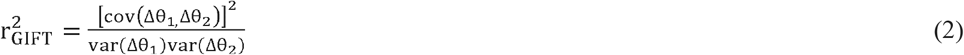

Where Δθ_1_ and Δθ_2_ represent two genetic paths with, var 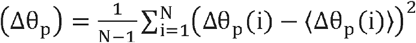 where p∈{1,2}and, 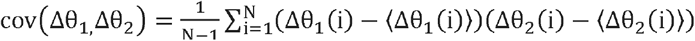. Note that with equation (2) the initial and arbitrary assignment of microstates is not fundamental. This can be demonstrated trivially as swapping ‘+1’ and ‘−1’ in Δθ_1_ (i) transforms it in −Δθ_1_ (i) with a resulting 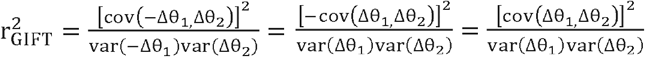.

Using equation (2), Fig.2D presents a new heatmap of SNP pairwise correlations for HMGA2. This result reveals stronger correlations between SNPs when genetic paths are considered that are not apparent in Fig.2B. Notably, examining the microstate composition of the two types of genetic paths in Fig.2E clarifies why they can be distinguished in Fig.2C, given the possible reassignment of microstates ‘+1’ and ‘−1’.

### GIFT’s determination of similarity between the genetic paths of significant SNPs across different chromosomes: Congruent genetic paths

Returning to the set of significant SNPs identified by GIFT in Table 2 (95% Bonferroni correction), heatmaps using equation (1) (Fig.3A) and equation (2) (Fig.3B) were generated to assess pairwise correlations between significant SNPs. In Figs.3A and 3B, the switch in colour for the SNP-Id is used represent the location of SNPs on different chromosomes. Notably, equation (1) does not suggest strong correlations between SNPs across different chromosomes (Fig.3A), whereas equation (2) indicates the opposite (Fig.3B). Since all SNPs represented in Fig.3B are significant, the absence of correlation between two SNPs implies that their genetic paths, i.e., their Δθ curves, differ in nature, for instance one following a parabolic trajectory and the other a sigmoidal one as exemplified by Figs.1C and 1D.

**Figure 3:**
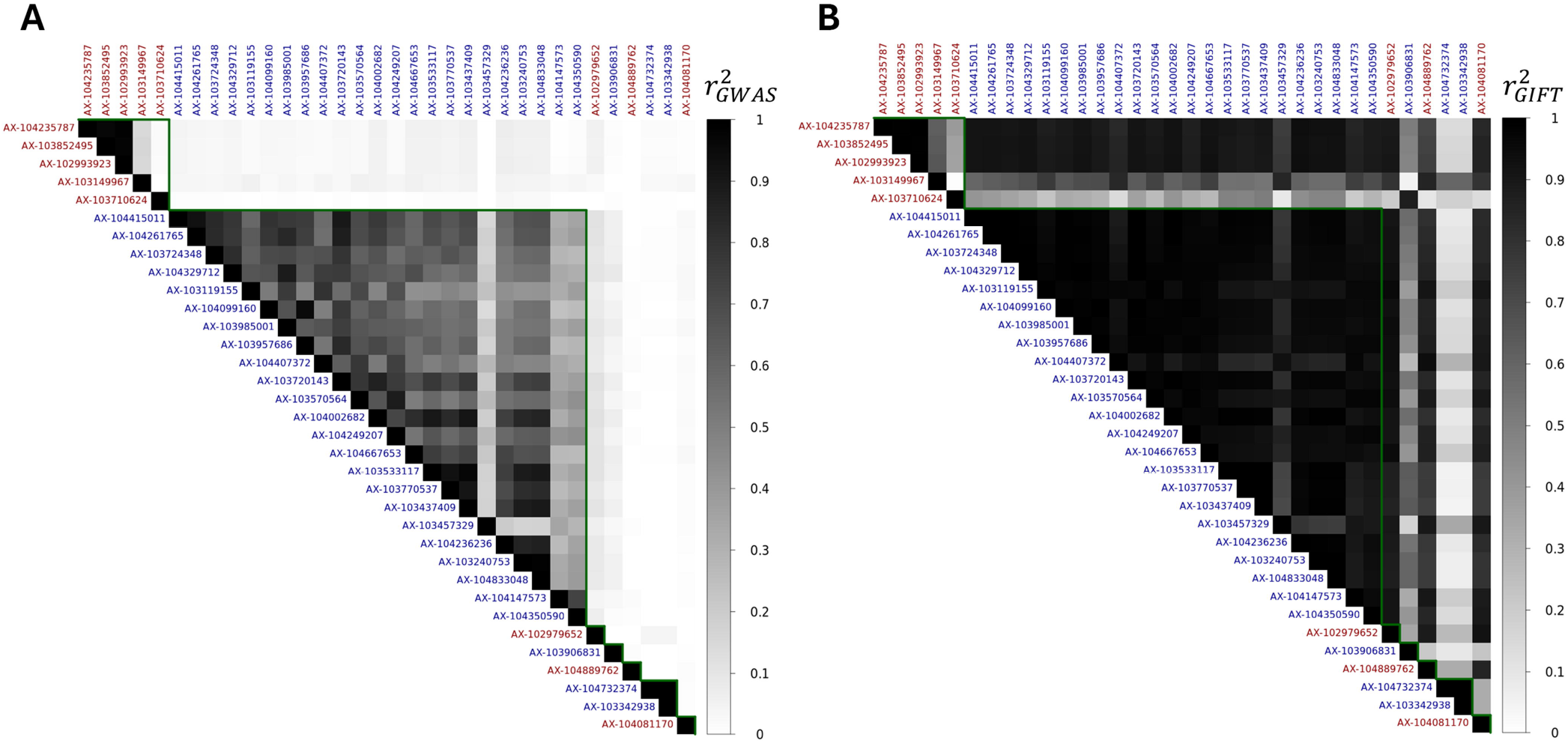
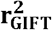 measure of SNP pairwise correlations to determine local gene networks. **(A)** 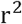-measure of pairwise correlation between significant SNPs (Table 2). The alternate red and blue colors of SNP Ids along the diagonal reflects that SNPs are located on different chromosomes. The green triangular line delineates SNP-Ids (Table 2) associated with specific chromosomes. **(B)** 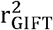-measure of pairwise correlations between SNPs (Table 2). Supplemental_Data_S5.xls provides the correlation values computed.

### GIFT’s determination of gene networks from congruent genetic paths

The key result presented in Fig.3 demonstrates the possibility of extracting gene network information from pairwise correlations between SNPs using equation (2), highlighting the emergence of potential networks based on similarities in the patterns (congruence) of two different SNP-related Δθs.

To assess the presence of such networks, FDR-adjusted p-values using the Benjamini-Hochberg process were used through which SNPs with a p-value<3% (Fig.1F) were selected, returning a total of 895 significant SNPs corresponding to 209 gene names. To extract strongly correlated SNPs using equation (2) a threshold of 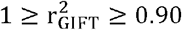 was set. As shown in Fig.4A, this process suggests strong correlations across chromosomes. To determine pairwise correlations between gene names, the average of pairwise correlations between SNPs pertaining to any two genes and noted 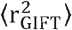 was used as numerical representative of pairwise correlation between genes.

**Figure 4.**
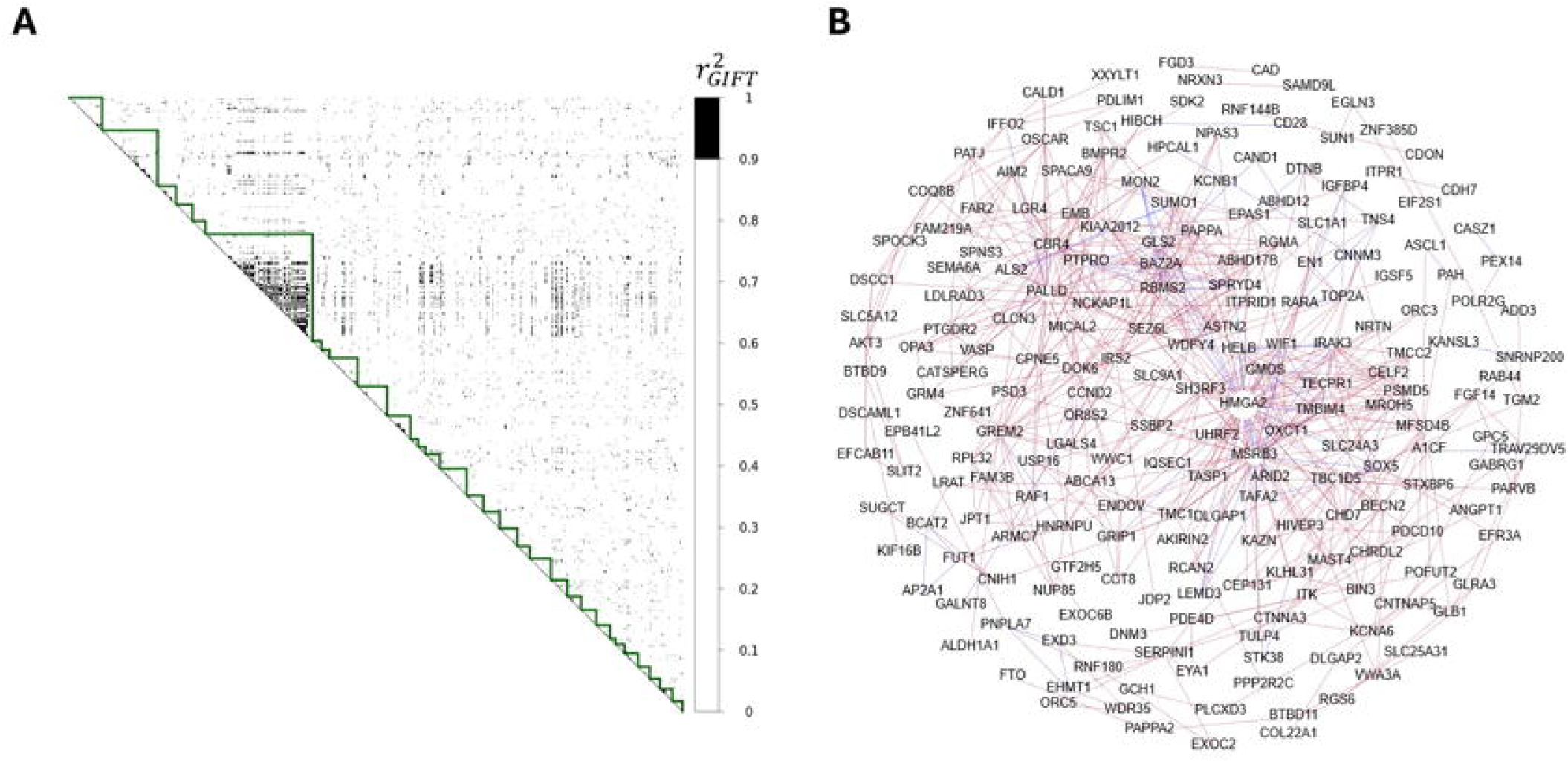
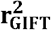 measure of gene names pairwise correlations to determine the isomorphic gene networks associated with “height at withers”. **(A)** Binary map of pairwise correlations between SNPs using GIFT, the green triangles separate the chromosomes. All chromosomes are present but in different proportions. **(B)** Resulting network of gene names associated with (A), where blue edges represent connections within the same chromosome, while red edges represent connections across different chromosomes. Supplemental_Data_S6.xls provides the average and variance of correlation values computed between genes noted 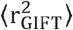.

Next, using gene names and a mean correlation value for correlations also within the range 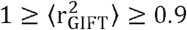, the *N* × *N* adjacency matrix defined as ***A*** = (*A*_*i,j*_), where *N* = 202 corresponds to known gene names, *A*_*i,j*_ = 1 if genes ‘i’ and ‘j’ are connected or zero otherwise, was computed and the gene networks visualized using the Fruchterman-Reingold layout in Gephi (30). With this layout genes are represented with their level of connectivity shown by a circle whose radius increases as the level of connectivity increases. As shown in Fig.4B, this representation differentiates genes with a high number of connections, which are typically positioned near the center of the network, from those with fewer connections, located at the periphery highlighting, in turn, a distinction between core and peripheral genes.

Fundamental to gene networks is the notion of gene centrality that determines a measure of a gene’s importance or influence within the network. One way to quantify centrality is through eigenvector centrality using the adjacency matrix previously computed. As the gene networks from Fig.4B are undirected, the matrix ***A*** is symmetric for each network, and the centrality measure for the gene ‘i’, denoted *g*_*i*_, is related to the centralities of i’s neighboring genes in the network as follows:

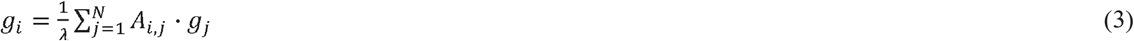

where λ is the largest eigenvalue of matrix ***A***. Defining the vector of centralities as 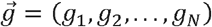,Eq.3 may be rewritten in matrix form:

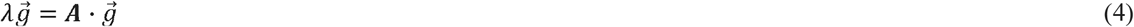

The vector 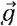 may be normalized so that the most important gene in the network has a weight of 1, that is, *max*(*g*_*i*_) = 1. Table 3 provides the name of central genes as well as their centrality measures found using equation (4).

**Table 3.** Centrality measures of isomorphic networks.

| Gene name | Eigenvector<br>rank value |
| --- | --- |
| HMGA2 | 1.0000 |
| MSRB3 (methionine sulfoxide reductase B3) | 0.6320 |
| SEZ6L (seizure related 6 homolog like) | 0.4686 |
| TMBIM4 (transmembrane BAX inhibitor motif-containing protein 4) | 0.4533 |
| TECPR1 (tectonin beta-Propeller repeat-containing protein 1) | 0.4460 |
| BAZ2A (bromodomain adjacent to zinc finger domain 2A) | 0.4316 |
| RBMS2 (RNA binding motif single stranded interacting protein 2) | 0.4316 |
| WIF1 (Wnt inhibitory factor 1) | 0.4255 |
| CELF2 (CUGBP Elav-like family member 2) | 0.4210 |
| WDFY4 (WD repeat- and FYVE domain-containing protein 4) | 0.4193 |

With the exceptions of HMGA2 and TMBIM4, the central genes identified are different from those listed in Tables 1 and 2. Additionally, it is visible from Table 3 that, except for HMGA2 and MSRB3, the centrality measures of genes drop rapidly to values ~0.4. Cross-referencing the second most central gene, MSRB3, using “body height” as a search term in the GWAS Catalog (29) confirmed that it is significantly associated with “body height” in humans (1). Additionally, MSRB3 was also cross-referenced in PubMed to determine whether its function is related to physiological processes associated with insulin action, insulin secretion, glucose uptake and homeostasis, energy metabolism, and metabolic disorders such as diabetes. In this context it was found that MSRB3, an endoplasmic reticulum (ER)-resident enzyme that protects cells from oxidative and ER stress by reducing oxidized methionine residues, has been implicated in the pathogenesis of insulin resistance and diabetes. A study using MSRB3 knockout mice showed that the absence of this enzyme inhibits the development of high-fat diet–induced insulin resistance. This effect is associated with increased levels of mitochondrial oxidative phosphorylation proteins, which promote mitochondrial biogenesis. These findings suggest that MSRB3 deficiency triggers compensatory mechanisms that preserves insulin sensitivity under metabolic stress (55).

## DISCUSSION

As statistical methods, GWAS and GIFT enable the determination of genotype-phenotype associations. The present study aimed to compare GWAS and GIFT results in identifying genes significantly associated with the trait “height at withers” in ponies, while simultaneously exploring potential links between this trait and equine metabolic syndrome (EMS), in a manner similar to the study by Norton et al., which was conducted on a larger sample size of 236 ponies (11).

Taken together, the results obtained: (i) confirm that GIFT extracts significantly more genetic information than GWAS, while also corroborating GWAS finding (HMGA2); and (ii) suggest that the genes identified by GIFT are strongly implicated in metabolic dysregulation involving insulin-related signaling pathways. Although the absence of SNPs associated with the insulin gene in either the GIFT or GWAS results may appear unexpected, this can be explained by the gene array used, which lacked coverage of the genomic regions corresponding to the insulin gene. Additionally, making use of GIFT’s framework, LD was redefined demonstrating the possibility to extract gene networks upon which centrality measures can be defined demonstrating that HMGA2 is indeed a central gene for the trait “height at withers”. Whilst this marks the first application of GIFT in equids with potential relevance to EMS, GIFT raises questions in relation to how genetic data must be interpreted.

The specific representation of scientific datasets shapes the foundation and structure of scientific knowledge bases. However, when the representation shifts, the underlying scientific concepts must also be redefined. When DDFs are used, their statistical summaries often based on their leading-order moments such as averages and variances, are expected to capture key scientific insights. In genetics, these statistical measures are closely tied to quantifiable concepts such as gene effect size and dominance for the average, and heritability for the variance. However, as the conceptual framework introduced by GIFT establishes the genetic path as a fundamental entity, one that cannot be reduced to conventional statistical summaries, this shift has important implications. Indeed, with GIFT a SNP can be significantly associated with a phenotype even when its effect size cannot be defined (see Fig.1D). Accordingly, if heredity is defined solely through variance, which itself is determined in relation to a mean or effect size, how can heredity be conceptualized for a significant SNP that lacks a measurable effect size (see Fig.1D)? These observations do not dispute the potential for any SNP to be associated with a trait or to exhibit heritability. Instead, they highlight the need to consider SNPs collectively rather than in isolation, positioning them within intricate dependencies and higher-order relationships that may elude traditional variance- or average-based approaches.

Traditionally, GWAS relies on genotyping a subset of SNPs, known as tag SNPs, which are in strong LD with many nearby SNPs. When a tag SNP shows a strong association with a trait, it suggests that a variant in LD with it may be associated with the trait. However, LD is typically determined based on bulk allele (or microstate) frequency differences calculated across the entire population. By using the detailed genetic paths provided by GIFT, it becomes now possible to refine the correlations between adjacent SNPs along a given chromosome, thereby offering a new perspective on LD. Note that, given the comprehensive information provided by GIFT, no base pair extensions were applied around tag SNPs in this present work. Due to the strength of correlations captured by GIFT, the method was extended to detect associations between SNPs across different chromosomes. Accordingly, the results demonstrate that the emergence of high-order congruent relationships become visible when the shape of genetic paths is considered (see Figs.2C,2D,3B,4A) demonstrating, in turn, that the genome is far more organized than initially thought.

GIFT’s ability to enhance the investigative power of datasets, in turn, enables discussion of a recently proposed concept in genetics: the omnigenic paradigm. The omnigenic paradigm for complex traits in genetics is a model that suggests that most genes in the genome contribute, directly or indirectly, to complex traits. This concept was first proposed by Boyle *et al*. and challenges the traditional polygenic model based on GWAS, which focuses on a limited set of key genes that cumulatively influence phenotypic trait (28). The key concepts of the omnigenic paradigm suggest that a small subset of core genes directly regulates the complex trait, while peripheral genes influence it indirectly through interactions with core genes within regulatory networks. While the theoretical and analytical frameworks, along with the present results from GIFT, support an omnigenic-like representation of complex traits, where an increasing number of SNPs are associated, it is important to acknowledge certain limitations. GIFT reinforces the concept of core versus peripheral genes, as illustrated in Fig.4B. However, contrary to the initial assumption that *most* genes in the genome contribute directly or indirectly to complex traits, GIFT suggests that only statistically significant genes are involved. In other words, it is unlikely that *most* genes contribute by default. In this context, GIFT can be seen as a hybrid conceptual framework for complex traits positioned between GWAS and the omnigenic paradigm.

Finally, it is essential to acknowledge the limitations of the present study:

i. While GIFT is a powerful tool for identifying genes associated with complex traits, rigorous validation of the results presented here requires an independent cohort, as a single association study cannot establish causality. Rather, such analyses identify candidate genes that may influence the trait. Furthermore, genes identified as significant require functional validation. In the absence of these additional steps, the reported associations should be considered putative.
ii. The trait selected for this study (“height at withers”) is highly heritable; therefore, the full potential of GIFT will likely become more apparent when applied to less heritable complex traits or diseases.

## DATA AVAILABILITY

Source data for this study are not publicly available due to commercial privacy or ethical restrictions. However, the source data are available to verified researchers upon request following instructions as explained in (25).

## SUPPLEMENTAL MATERIAL

All supplementary materials are available at: https://doi.org/10.6084/m9.figshare.32005926

## ACKNOWLEDGMENTS

We would like to thank Dr Oswald Matika and Dr Sian Bray for useful discussions concerning data analysis.

## AUTHOR CONTRIBUTIONS

CR conceptualized GIFT; CR, JW formalized GIFT; PK coded GIFT simulations; SG computed network centrality results; the dataset used was designed and obtained by EK, PH, NMG, JE; paper was written by CR; paper was proofread by EK, PH, NMG, JE, AP, CR.

## REFERENCES

1. Yengo L, Vedantam S, Marouli E, Sidorenko J, Bartell E, Sakaue S, Graff M, Eliasen AU, Jiang Y, Raghavan S, Miao J, Arias JD, Graham SE, Mukamel RE, Spracklen CN, Yin X, Chen S-H, Ferreira T, Highland HH, Ji Y, Karaderi T, Lin K, Lüll K, Malden DE, Medina-Gomez C, Machado M, Moore A, Rüeger S, Sim X, Vrieze S, Ahluwalia TS, Akiyama M, Allison MA, Alvarez M, Andersen MK, Ani A, Appadurai V, Arbeeva L, Bhaskar S, Bielak LF, Bollepalli S, Bonnycastle LL, Bork-Jensen J, Bradfield JP, Bradford Y, Braund PS, Brody JA, Burgdorf KS, Cade BE, Cai H, Cai Q, Campbell A, Cañadas-Garre M, Catamo E, Chai J-F, Chai X, Chang L-C, Chang Y-C, Chen C-H, Chesi A, Choi SH, Chung R-H, Cocca M, Concas MP, Couture C, Cuellar-Partida G, Danning R, Daw EW, Degenhard F, Delgado GE, Delitala A, Demirkan A, Deng X, Devineni P, Dietl A, Dimitriou M, Dimitrov L, Dorajoo R, Ekici AB, Engmann JE, Fairhurst-Hunter Z, Farmaki A-E, Faul JD, Fernandez-Lopez J-C, Forer L, Francescatto M, Freitag-Wolf S, Fuchsberger C, Galesloot TE, Gao Y, Gao Z, Geller F, Giannakopoulou O, Giulianini F, Gjesing AP, Goel A, Gordon SD, Gorski M, Grove J, Guo X, Gustafsson S, Haessler J, Hansen TF, Havulinna AS, Haworth SJ, He J, Heard-Costa N, Hebbar P, Hindy G, Ho Y-LA, Hofer E, Holliday E, Horn K, Hornsby WE, Hottenga J-J, Huang H, Huang J, Huerta-Chagoya A, Huffman JE, Hung Y-J, Huo S, Hwang MY, Iha H, Ikeda DD, Isono M, Jackson AU, Jäger S, Jansen IE, Johansson I, Jonas JB, Jonsson A, Jørgensen T, Kalafati I-P, Kanai M, Kanoni S, Kårhus LL, Kasturiratne A, Katsuya T, Kawaguchi T, Kember RL, Kentistou KA, Kim H-N, Kim YJ, Kleber ME, Knol MJ, Kurbasic A, Lauzon M, Le P, Lea R, Lee J-Y, Leonard HL, Li SA, Li X, Li X, Liang J, Lin H, Lin S-Y, Liu J, Liu X, Lo KS, Long J, Lores-Motta L, Luan J, Lyssenko V, Lyytikäinen L-P, Mahajan A, Mamakou V, Mangino M, Manichaikul A, Marten J, Mattheisen M, Mavarani L, McDaid AF, Meidtner K, Melendez TL, Mercader JM, Milaneschi Y, Miller JE, Millwood IY, Mishra PP, Mitchell RE, Møllehave LT, Morgan A, Mucha S, Munz M, Nakatochi M, Nelson CP, Nethander M, Nho CW, Nielsen AA, Nolte IM, Nongmaithem SS, Noordam R, Ntalla I, Nutile T, Pandit A, Christofidou P, Pärna K, Pauper M, Petersen ERB, Petersen LV, Pitkänen N, Polašek O, Poveda A, Preuss MH, Pyarajan S, Raffield LM, Rakugi H, Ramirez J, Rasheed A, Raven D, Rayner NW, Riveros C, Rohde R, Ruggiero D, Ruotsalainen SE, Ryan KA, Sabater-Lleal M, Saxena R, Scholz M, Sendamarai A, Shen B, Shi J, Shin JH, Sidore C, Sitlani CM, Slieker RC, Smit RAJ, Smith AV, Smith JA, Smyth LJ, Southam L, Steinthorsdottir V, Sun L, Takeuchi F, Tallapragada DSP, Taylor KD, Tayo BO, Tcheandjieu C, Terzikhan N, Tesolin P, Teumer A, Theusch E, Thompson DJ, Thorleifsson G, Timmers PRHJ, Trompet S, Turman C, Vaccargiu S, van der Laan SW, van der Most PJ, van Klinken JB, van Setten J, Verma SS, Verweij N, Veturi Y, Wang CA, Wang C, Wang L, Wang Z, Warren HR, Bin Wei W, Wickremasinghe AR, Wielscher M, Wiggins KL, Winsvold BS, Wong A, Wu Y, Wuttke M, Xia R, Xie T, Yamamoto K, Yang J, Yao J, Young H, Yousri NA, Yu L, Zeng L, Zhang W, Zhang X, Zhao J-H, Zhao W, Zhou W, Zimmermann ME, Zoledziewska M, Adair LS, Adams HHH, Aguilar-Salinas CA, Al-Mulla F, Arnett DK, Asselbergs FW, Åsvold BO, Attia J, Banas B, Bandinelli S, Bennett DA, Bergler T, Bharadwaj D, Biino G, Bisgaard H. A saturated map of common genetic variants associated with human height. Nature 610: 704–712, 2022. doi: 10.1038/s41586-022-05275-y.

2. Steiner CC, Putnam AS, Hoeck PEA, Ryder OA. Conservation genomics of threatened animal species. Annu Rev Anim Biosci 1: 261–281, 2013. doi: 10.1146/annurev-animal-031412-103636.

3. Rauch C, Wattis J, Bray S. On the Meaning of Averages in Genome-wide Association Studies: What Should Come Next? Organisms Journal of Biological Sciences 6: 7–22, 2023. doi: 10.13133/2532-5876/17811.

4. Nelson RM, Pettersson ME, Carlborg Ö. A century after Fisher: time for a new paradigm in quantitative genetics. Trends in geneticsflJ: TIG 29: 669–676, 2013. doi: 10.1016/j.tig.2013.09.006.

5. Wattis JAD, Bray SM, Kyratzi P, Rauch C. Analysis of phenotype-genotype associations using genomic informational field theory (GIFT). J Theor Biol 548: 111198, 2022. doi: 10.1016/j.jtbi.2022.111198.

6. Rauch C, Kyratzi P, Blott S, Bray S, Wattis J. GIFT: new method for the genetic analysis of small gene effects involving small sample sizes. Phys Biol 20, 2022. doi: 10.1088/1478-3975/ac99b3.

7. Kyratzi P, Matika O, Brassington AH, Clare CE, Xu J, Barrett DA, Emes RD, Archibald AL, Paldi A, Sinclair KD, Wattis J, Rauch C. Investigative power of genomic informational field theory relative to genomewide association studies for genotype-phenotype mapping. Physiological Genomics 56: 791–806, 2024. doi: 10.1152/physiolgenomics.00049.2024.

8. Sadek MH, Al-Aboud AZ, Ashmawy AA. Factor analysis of body measurements in Arabian horses. J Anim Breed Genet 123: 369–377, 2006. doi: 10.1111/j.1439-0388.2006.00618.x.

9. Molina A, Valera M, Santos RD, Rodero A. Genetic parameters of morphofunctional traits in Andalusian horse. Livestock Production Science 60: 295–303, 1999. doi: 10.1016/S0301-6226(99)00101-3.

10. Outram AK, Stear NA, Bendrey R, Olsen S, Kasparov A, Zaibert V, Thorpe N, Evershed RP. The earliest horse harnessing and milking. Science 323: 1332–1335, 2009. doi: 10.1126/science.1168594.

11. Norton EM, Avila F, Schultz NE, Mickelson JR, Geor RJ, McCue ME. Evaluation of an HMGA2 variant for pleiotropic effects on height and metabolic traits in ponies. J Vet Intern Med 33: 942–952, 2019. doi: 10.1111/jvim.15403.

12. Krebs LC, Santos MM de M, Siqueira MC, Araujo BPG de, Solar Diaz IDP, Costa RB, Oliveira CA de A, Barbero MMD, de Camargo GMF, Godoi FN de. Candidate genes for height measurements in Campolina horses [Online]. Anim Prod Sci 64, 2024. 10.1071/AN23071.

13. Junior AB, Quirino CR, Vega WHO, Rua MAS, David CMG, Jardim JG. Polymorphisms in the LASP1 gene allow selection for smaller stature in ponies. Livestock Science 216: 160–164, 2018. doi: 10.1016/j.livsci.2018.07.015.

14. Makvandi-Nejad S, Hoffman GE, Allen JJ, Chu E, Gu E, Chandler AM, Loredo AI, Bellone RR, Mezey JG, Brooks SA, Sutter NB. Four loci explain 83% of size variation in the horse. PLoS One 7: e39929, 2012. doi: 10.1371/journal.pone.0039929.

15. Frischknecht M, Jagannathan V, Plattet P, Neuditschko M, Signer-Hasler H, Bachmann I, Pacholewska A, Drögemüller C, Dietschi E, Flury C, Rieder S, Leeb T. A Non-Synonymous HMGA2 Variant Decreases Height in Shetland Ponies and Other Small Horses. PLoS One 10: e0140749, 2015. doi: 10.1371/journal.pone.0140749.

16. Norton EM, Schultz NE, Rendahl AK, Mcfarlane D, Geor RJ, Mickelson JR, McCue ME. Heritability of metabolic traits associated with equine metabolic syndrome in Welsh ponies and Morgan horses. Equine Vet J 51: 475–480, 2019. doi: 10.1111/evj.13053.

17. Tozaki T, Kikuchi M, Kakoi H, Hirota K-I, Nagata S-I. A genome-wide association study for body weight in Japanese Thoroughbred racehorses clarifies candidate regions on chromosomes 3, 9, 15, and 18. J Equine Sci 28: 127–134, 2017. doi: 10.1294/jes.28.127.

18. Metzger J, Schrimpf R, Philipp U, Distl O. Expression levels of LCORL are associated with body size in horses. PLoS One 8: e56497, 2013. doi: 10.1371/journal.pone.0056497.

19. Reich P, Möller S, Stock KF, Nolte W, von Depka Prondzinski M, Reents R, Kalm E, Kühn C, Thaller G, Falker-Gieske C, Tetens J. Genomic analyses of withers height and linear conformation traits in German Warmblood horses using imputed sequence-level genotypes. Genet Sel Evol 56: 45, 2024. doi: 10.1186/s12711-024-00914-6.

20. Signer-Hasler H, Flury C, Haase B, Burger D, Simianer H, Leeb T, Rieder S. A genome-wide association study reveals loci influencing height and other conformation traits in horses. PLoS One 7: e37282, 2012. doi: 10.1371/journal.pone.0037282.

21. Tetens J, Widmann P, Kühn C, Thaller G. A genome-wide association study indicates LCORL/NCAPG as a candidate locus for withers height in German Warmblood horses. Anim Genet 44: 467–471, 2013. doi: 10.1111/age.12031.

22. Staiger EA, Al Abri MA, Pflug KM, Kalla SE, Ainsworth DM, Miller D, Raudsepp T, Sutter NB, Brooks SA. Skeletal variation in Tennessee Walking Horses maps to the LCORL/NCAPG gene region. Physiol Genomics 48: 325–335, 2016. doi: 10.1152/physiolgenomics.00100.2015.

23. Suontama M, Saastamoinen MT, Ojala M. Estimates of non-genetic effects and genetic parameters for body measures and subjectively scored traits in Finnhorse trotters. Livestock Science 124: 205–209, 2009. doi: 10.1016/j.livsci.2009.01.017.

24. Zechner P, Zohman F, Sölkner J, Bodo I, Habe F, Marti E, Brem G. Morphological description of the Lipizzan horse population. Livestock Production Science 69: 163–177, 2001. doi: 10.1016/S0301-6226(00)00254-2.

25. Knowles EJ, Elliott J, Harris PA, Chang Y-M, Menzies-Gow NJ. Predictors of laminitis development in a cohort of nonlaminitic ponies. Equine Vet J 55: 12–23, 2023. doi: 10.1111/evj.13572.

26. Clark BL, Bamford NJ, Stewart AJ, McCue ME, Rendahl A, Bailey SR, Bertin F-R, Norton EM. Evaluation of an HMGA2 variant contribution to height and basal insulin concentrations in ponies. J Vet Intern Med 37: 1186–1192, 2023. doi: 10.1111/jvim.16723.

27. Gadsby S, Rauch C, Wattis JAD. Identification of significant SNPs and the quantification of correlation using genomic informational field theory (GIFT). Mathematical Biosciences 393: 109606, 2026. doi: 10.1016/j.mbs.2025.109606.

28. Boyle EA, Li YI, Pritchard JK. An Expanded View of Complex Traits: From Polygenic to Omnigenic. Cell 169: 1177–1186, 2017. doi: 10.1016/j.cell.2017.05.038.

29. Cerezo M, Sollis E, Ji Y, Lewis E, Abid A, Bircan KO, Hall P, Hayhurst J, John S, Mosaku A, Ramachandran S, Foreman A, Ibrahim A, McLaughlin J, Pendlington Z, Stefancsik R, Lambert SA, McMahon A, Morales J, Keane T, Inouye M, Parkinson H, Harris LW. The NHGRI-EBI GWAS Catalog: standards for reusability, sustainability and diversity. Nucleic Acids Research 53: D998–D1005, 2025. doi: 10.1093/nar/gkae1070.

30. Bastian M, Heymann S, Jacomy M. Gephi: An Open Source Software for Exploring and Manipulating Networks. ICWSM 3: 361–362, 2009. doi: 10.1609/icwsm.v3i1.13937.

31. Horiba N, Masuda S, Takeuchi A, Takeuchi D, Okuda M, Inui K. Cloning and characterization of a novel Na+-dependent glucose transporter (NaGLT1) in rat kidney. J Biol Chem 278: 14669–14676, 2003. doi: 10.1074/jbc.M212240200.

32. UniProt: the Universal Protein Knowledgebase in 2025. Nucleic Acids Res 53: D609–D617, 2025. doi: 10.1093/nar/gkae1010.

33. Szklarczyk D, Kirsch R, Koutrouli M, Nastou K, Mehryary F, Hachilif R, Gable AL, Fang T, Doncheva NT, Pyysalo S, Bork P, Jensen LJ, von Mering C. The STRING database in 2023: protein-protein association networks and functional enrichment analyses for any sequenced genome of interest. Nucleic Acids Res 51: D638–D646, 2023. doi: 10.1093/nar/gkac1000.

34. Naval-Sánchez M, Porto-Neto LR, Cardoso DF, Hayes BJ, Daetwyler HD, Kijas J, Reverter A. Selection signatures in tropical cattle are enriched for promoter and coding regions and reveal missense mutations in the damage response gene HELB. Genet Sel Evol 52: 27, 2020. doi: 10.1186/s12711-020-00546-6.

35. Webster MT, Kamgari N, Perloski M, Hoeppner MP, Axelsson E, Hedhammar Å, Pielberg G, Lindblad-Toh K. Linked genetic variants on chromosome 10 control ear morphology and body mass among dog breeds. BMC Genomics 16: 474, 2015. doi: 10.1186/s12864-015-1702-2.

36. Posbergh CJ, Huson HJ. All sheeps and sizes: a genetic investigation of mature body size across sheep breeds reveals a polygenic nature. Anim Genet 52: 99–107, 2021. doi: 10.1111/age.13016.

37. Su L, Deng Z, Leng F. The Mammalian High Mobility Group Protein AT-Hook 2 (HMGA2): Biochemical and Biophysical Properties, and Its Association with Adipogenesis. Int J Mol Sci 21, 2020. doi: 10.3390/ijms21103710.

38. Banga A, Unal R, Tripathi P, Pokrovskaya I, Owens RJ, Kern PA, Ranganathan G. Adiponectin translation is increased by the PPARγ agonists pioglitazone and ω-3 fatty acids. American Journal of Physiology-Endocrinology and Metabolism 296: E480–E489, 2009. doi: 10.1152/ajpendo.90892.2008.

39. Seyyed Amin Seyyed Rezaei, Akbar Amirfiroozi, Moein Kohkalani, Maghsoud Mehri. Transcription Factors in Type 2 Diabetes: Molecular Mechanisms of Insulin Resistance and β-Cell Dysfunction. Current Diabetes Reviews 22: 1–15, 2026. doi: 10.2174/0115733998443779260129064444.

40. Sokolowski EK, Kursawe R, Selvam V, Bhuiyan RM, Thibodeau A, Zhao C, Spracklen CN, Ucar D, Stitzel ML. Multi-omic human pancreatic islet endoplasmic reticulum and cytokine stress response mapping provides type 2 diabetes genetic insights. Cell Metabolism 36: 2468-2488.e7, 2024. doi: 10.1016/j.cmet.2024.09.006.

41. Raab S, Gadault A, Very N, Decourcelle A, Baldini S, Schulz C, Mortuaire M, Lemaire Q, Hardivillé S, Dehennaut V, El Yazidi-Belkoura I, Vercoutter-Edouart A-S, Panasyuk G, Lefebvre T. Dual regulation of fatty acid synthase (FASN) expression by O-GlcNAc transferase (OGT) and mTOR pathway in proliferating liver cancer cells. Cellular and Molecular Life Sciences 78: 5397–5413, 2021. doi: 10.1007/s00018-021-03857-z.

42. Wang H, Luo QF, Peng AF, Long XH, Wang TF, Liu ZL, Zhang GM, Zhou RP, Gao S, Zhou Y, Chen WZ. Positive feedback regulation between Akt phosphorylation and fatty acid synthase expression in osteosarcoma. Int J Mol Med 33: 633–639, 2014. doi: 10.3892/ijmm.2013.1602.

43. Zhang J, Chen T, Wu W, Hu C, Wang B, Jia X, Ye M. Carbonyl reductase 4 suppresses colorectal cancer progression through the DNMT3B/CBR4/FASN/mTOR axis. Cancer Cell Int 25: 146, 2025. doi: 10.1186/s12935-025-03776-0.

44. Jin L. The actin associated protein palladin in smooth muscle and in the development of diseases of the cardiovasculature and in cancer. J Muscle Res Cell Motil 32: 7–17, 2011. doi: 10.1007/s10974-011-9246-9.

45. Tsakiridis T, Tong P, Matthews B, Tsiani E, Bilan PJ, Klip A, Downey GP. Role of the actin cytoskeleton in insulin action. Microsc Res Tech 47: 79–92, 1999. doi: 10.1002/(SICI)1097-0029(19991015)47:2<79::AID-JEMT1>3.0.CO;2-S.

46. Huang S-MA, Hancock MK, Pitman JL, Orth AP, Gekakis N. Negative regulators of insulin signaling revealed in a genome-wide functional screen. PLoS One 4: e6871, 2009. doi: 10.1371/journal.pone.0006871.

47. Xie H, Liu X, Li S, Wang M, Li Y, Chen T, Li L, Wang F, Xiao X. Tissue adaptation to metabolic stress: insights from SUMOylation. Front Endocrinol (Lausanne) 15: 1434338, 2024. doi: 10.3389/fendo.2024.1434338.

48. Kamynina E, Stover PJ. The Roles of SUMO in Metabolic Regulation. Adv Exp Med Biol 963: 143–168, 2017. doi: 10.1007/978-3-319-50044-7_9.

49. Leonardini A, Laviola L, Perrini S, Natalicchio A, Giorgino F. Cross-Talk between PPARgamma and Insulin Signaling and Modulation of Insulin Sensitivity. PPAR Res 2009: 818945, 2009. doi: 10.1155/2009/818945.

50. Mikkonen L, Hirvonen J, Jänne OA. SUMO-1 Regulates Body Weight and Adipogenesis via PPARγ in Male and Female Mice. Endocrinology 154: 698–708, 2013. doi: 10.1210/en.2012-1846.

51. Chen P, Cheng H, Zheng F, Li S, Bornhorst J, Yang B, Lee KH, Ke T, Li Y, Schwerdtle T, Yang X, Bowman AB, Aschner M. BTBD9 attenuates manganese-induced oxidative stress and neurotoxicity by regulating insulin growth factor signaling pathway. Hum Mol Genet 31: 2207–2222, 2022. doi: 10.1093/hmg/ddac025.

52. Chen P, Zheng F, Li S, Cheng H, Bornhorst J, Li Y, Yang B, Lee KH, Ke T, Schwerdtle T, Yang X, Bowman AB, Aschner M. BTBD9 is a novel component of IGF signaling and regulates manganese-induced dopaminergic dysfunction.

53. Chen J, Meng Y, Zhou J, Zhuo M, Ling F, Zhang Y, Du H, Wang X. Identifying candidate genes for Type 2 Diabetes Mellitus and obesity through gene expression profiling in multiple tissues or cells. J Diabetes Res 2013: 970435, 2013. doi: 10.1155/2013/970435.

54. Hill WG, Robertson A. Linkage disequilibrium in finite populations. Theoretical and Applied Genetics 38: 226–231, 1968. doi: 10.1007/BF01245622.

55. Cha H-N, Woo C-H, Kim H-Y, Park S-Y. Methionine sulfoxide reductase B3 deficiency inhibits the development of diet-induced insulin resistance in mice. Redox Biology 38: 101823, 2021. doi: 10.1016/j.redox.2020.101823.

