## Supplemental Text S1 for "Genomic Informational Field Theory (GIFT) to identify genetic associations of a complex trait using a small sample size"

**Introduction to GIFT METHODOLOGY**

GIFT is a method designed to decouple the notion of precision in phenotypic measurements from that of sample size. Put differently, it provides genotype-phenotype associations without relying on data categorization or grouping. To develop the theoretical framework underpinning GIFT, it is crucial to understand what increasing precision in phenotypic measurements entails with current methods.

**Deconstructing GWAS to let GIFT emerge**

Consider a specific single nucleotide polymorphism (SNP) in the genome, assumed to be associated with a given phenotype in a Mendelian manner. We shall refer to this SNP as SNP1. With current GWAS methods, which are based on Fisher’s theory, the phenotype distribution density function (see below Fig.1A-top panel) is decomposed into the distribution density functions of distinct genetic microstates (Fig.1B-top panel). Using GWAS, the effect size (2a, Fig.1C-top panel) and dominance (d, Fig.1C-top panel) are derived via linear interpolation of averages (Fig.1C).

However, when phenotypic measurement precision increases without changing the sample size (fixed at 1,000 individuals in this simulation, Fig.1), the distribution density functions transform into barcodes (Fig.1A-bottom panel and Fig.1B-bottom panel). At the genetic level, Fig.1B illustrates that decoupling sample size and precision is equivalent to comprehending the overall configuration of the different microstates represented in the barcode (Fig.1B-bottom panel).

When examining the barcode of microstates, the first point to note is that the precise positioning of bars along the space of phenotypic values (i.e., their exact locations on the x-axis in Fig.1A-bottom panel) depends on the sampling process. Specifically, different individuals with slightly varying phenotypic values would result in different bar positions in the phenotypic value space. Therefore, the absolute positioning of the bars is not as important as their relative positioning that is, their order in the sequence they form.

Because the barcode in Fig.1B-bottom panel reflects the association of SNP1 with the phenotype, the different microstates are segregated. The resulting configuration of the string of microstates from Fig.1B-bottom panel can be represented as follows:

SNP1: [+1, +1, +1, +1, 0, +1, +1, …, +1, 0, 0, 0, -1, 0, -1, …, -1, 0, -1, -1, -1, -1, -1]	(1)

To capture the overall configuration of the string, one approach is to construct a new string, denoted as $\theta=\left[ \left\{ \theta\left( i \right) \right\} \right]$, with the same dimension as SNP1 that is, having the same number of components. In this string, the component at position $i$ results from the cumulative sum of microstates in SNP1 (Eq.1) up to position $i$. Specifically, $\theta\left( i=1 \right)=+1=1$, $\theta\left( i=2 \right)=+1+1=2$, $\theta\left( i=3 \right)=+1+1+1=3$, and so on until, $\theta\left( i=N \right)=N_{+}-N_{-}$, where $N_{+}$ and $N_{-}$ are the number of ‘+1’ and ‘-1’ states in SNP1, respectively. We shall call this type of cumulative sum a genetic path.

Now consider another SNP in the genome, denoted SNP2, which is not associated with the phenotype. For SNP2 the configuration of microstates should appear random, meaning the bars’ colors are not partitioned or segregated. An example of such a case could be:

SNP2: [-1, +1, 0, -1, -1, +1, +1, …, -1, +1, +1, 0, -1, 0, +1, …, 0, 0, -1, +1, 0, +1, -1]	(2)

The difference between (1) and (2), namely, the presence or absence of a genotype-phenotype association, lies in the concepts of ‘scrambling’ or ‘mixing’ of microstates. While SNP1 and SNP2 correspond to different genome positions and are not directly comparable, one could artificially scramble SNP1 via random permutations. In such a scrambled state, there would be no association between the genotype (SNP1) and the phenotype, for the phenotypic information used initially to order microstates would be lost. By calculating the cumulative sum of microstates for this scrambled configuration, one could determine a null hypothesis for the genetic path related to SNP1.

One issue to address, however, is that while SNP1 is unique, many different random configurations are possible through scrambling. To address this, one might ask what sort of average string would emerge after considering an infinite number of random permutations of SNP1. This is straightforward, as it is equivalent to determining the probability of each microstate’s presence in the string. If $N_{+}$, $N_{0}$ and $N_{-}$ represent the counts of ‘+1’, ‘0’ and ‘-1’ microstates in SNP1, the probability $p_{q}$ (where $q\in\left\{ +,0,- \right\}$) to find the microstate $q$ is, $p_{q}=N_{q}/N$, where $N=N_{+}$+$N_{0}$+$N_{-}$ is the total number of microstates. Each component of the string must then take the weighted average value, $+1\cdot p_{+}+0\cdot p_{0}-1\cdot p_{-}=p_{+}-p_{-}$.

Thus, denoting $\theta_{0}=\left[ \left\{ \theta_{0}\left( i \right) \right\} \right]$ as the new cumulative sum of components after considering an infinite number of random permutations, one concludes, $\theta_{0}\left( i \right)=\left( p_{+}-p_{-} \right)i$.

The two genetic paths, $\theta=\left[ \left\{ \theta\left( i \right) \right\} \right]$ and $\theta_{0}=\left[ \left\{ \left( p_{+}-p_{-} \right)i \right\} \right]$, can then be compared at every position using their difference:

$\Delta\theta=\theta-\theta_{0}=\left[ \left\{ \theta\left( i \right)-\left( p_{+}-p_{-} \right)i \right\} \right]$	(3)

Unlike GWAS, which determines associations using averages and linear interpolation (as illustrated by the green line in Fig.1D), GIFT focuses on assessing the significance of amplitude differences.

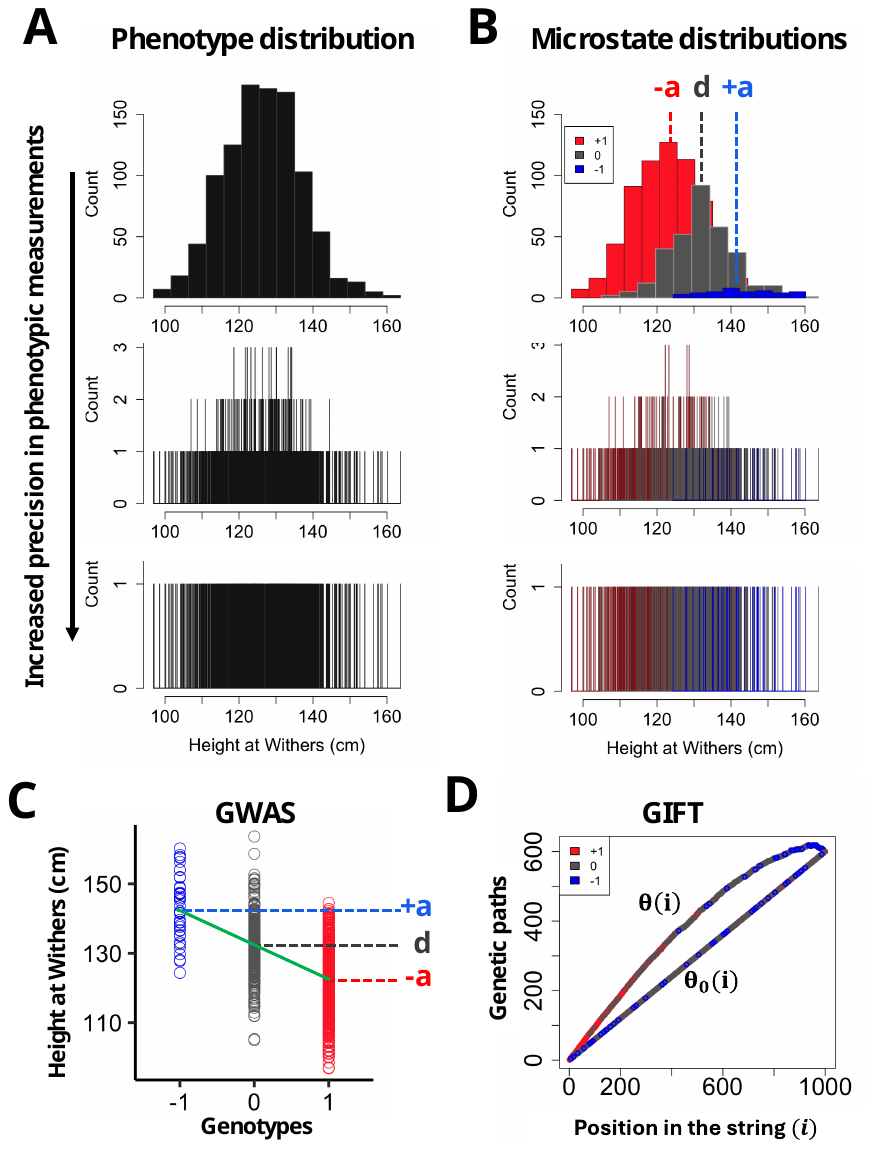

**Figure 1.** GIFT, a novel approach for genetic inferences. *(A)* & *(B)* In diploid organisms, a bi-allelic genetic marker (A or a) can exist in three genotypic states: ‘+1’ (aa, red), ‘0’ (Aa/aA, grey), and ‘-1’ (AA, blue). GWAS relies on distribution density functions formed by grouping data into bins. The phenotype distribution density (*(A)* top panel) is decomposed into the genetic microstate distributions (*(B)* top panel) for each SNP. *(C)* GWAS assesses SNP-phenotype associations using allelic dosage. A significant nonzero slope in the fitted regression line indicates an association. *(D)* Higher precision requires narrower bin widths, demanding larger sample sizes to maintain statistical power. To address this, GIFT deconstructs density functions, increasing phenotypic precision without altering sample size (*(A)*&*(B)* top-to-bottom panels). This transformation generates a barcode-like sequence of microstates, which GIFT analyzes via, $\Delta\theta=\theta-\theta_{0}$ (Eq. 3), measuring non-random microstate organization. Note: the simulation presented adheres to Fisher’s model with a constant sample size of 1000 and a normally distributed phenotype (mean: 132 cm, variance: 10 cm²). Each microstate follows a normal distribution with a gene/size effect (2a) equal to the phenotype’s standard deviation without dominance (d=0), and the genotype frequencies, aa (64%), Aa/aA (32%), and AA (4%), follow Hardy-Weinberg equilibrium.

**Discriminative and investigative powers of GIFT**

The new properties of GIFT can now be discussed. The first property is what may be termed the "discriminative power" of GIFT. Unlike Fisher’s method, which considers a null hypothesis at the population level, where overlapping microstate distributions make differences in averages impossible to determine (Fig. 1C), GIFT defines the null hypothesis at the SNP (genetic) level. This means that each SNP has its own null hypothesis, offering greater discriminative power compared to GWAS.

The second property pertains to GIFT's "investigative power." GIFT seeks to extract the non-random organization of microstates from $\Delta\theta$. Detecting some level of organization is also a goal of GWAS, which does so using averages. Indeed, GWAS evaluates whether a partitioning of microstates exists or is statistically significant, as clearly illustrated in Fig.1C. However, the key difference lies in the granularity of the organization that each method seeks to detect. The granularity is high with GWAS, which using data grouping (granules) examines variations through averages and variances, whereas GIFT focuses on capturing broader, non-random patterns of microstate organization, effectively bypassing granularity.

The distinction between the investigative powers of GWAS and GIFT can be best exemplified through an example. After pre-correcting the phenotype "height at withers" (see Methods in the main text) to work with residual phenotypic values, we present the results in Fig.2A, Fig.2B, and Fig.2C. These figures show, in the left panels, the phenotype allelic dosage using the GWAS method and, in the middle and right panels, the genetic paths derived using the GIFT method. The data correspond to three SNPs: AX-104329712 (Fig. 2A), AX-103149967 (Fig.2B), and AX-103149967 (Fig. 2C), located on Chromosomes 6 (position 66597482), 2 (position 82564015), and 1 (position 677336), respectively.

Focusing on the information extracted by GWAS, Fig.2A (left panel) resembles Fig.1C, obtained through simulation, where the asymmetrical partitioning of homozygote microstates, represented by their colors, reveals a certain level of organization. This, in turn, suggests an association between genotype and phenotype. In this case, a gene/size effect of $a=-2.11\cdot{10}^{-5} a.u.$ (a.u.: arbitrary unit), a dominance, $d=5.23\cdot{10}^{-6} a.u.$, and an $r^{2}$-value of $0.174$, can be determined with a corresponding, ${-Log}_{10}\left( p_{\mathrm{GWAS}} \right)=7.2$. By contrast, Fig.2B (left panel), shows that GWAS does not indicate any significant association (${-Log}_{10}\left( p_{\mathrm{GWAS}} \right)=0.5, a=-6.08\cdot{10}^{-6} a.u., d=5.23\cdot{10}^{-6}$ a.u., $r^{2}=0.017$).

Let us now examine the information extracted by GIFT. The middle panel of Fig.2A shows genetic paths, similar to those in Fig.1D. The shape for $\Delta\theta$ (Fig.2A, right panel) is parabolic, indicating a non-random distribution of microstates. Similarly, the comparison between $\theta$ and $\theta_{0}$ in Fig.2B (middle panel) suggests a non-random organization of microstates, which is further supported by the sigmoidal curve observed for underlined $\Delta\theta$ (Fig.2B, right panel). In both cases, GIFT indicates that these genetic paths, although distinct, are not random.

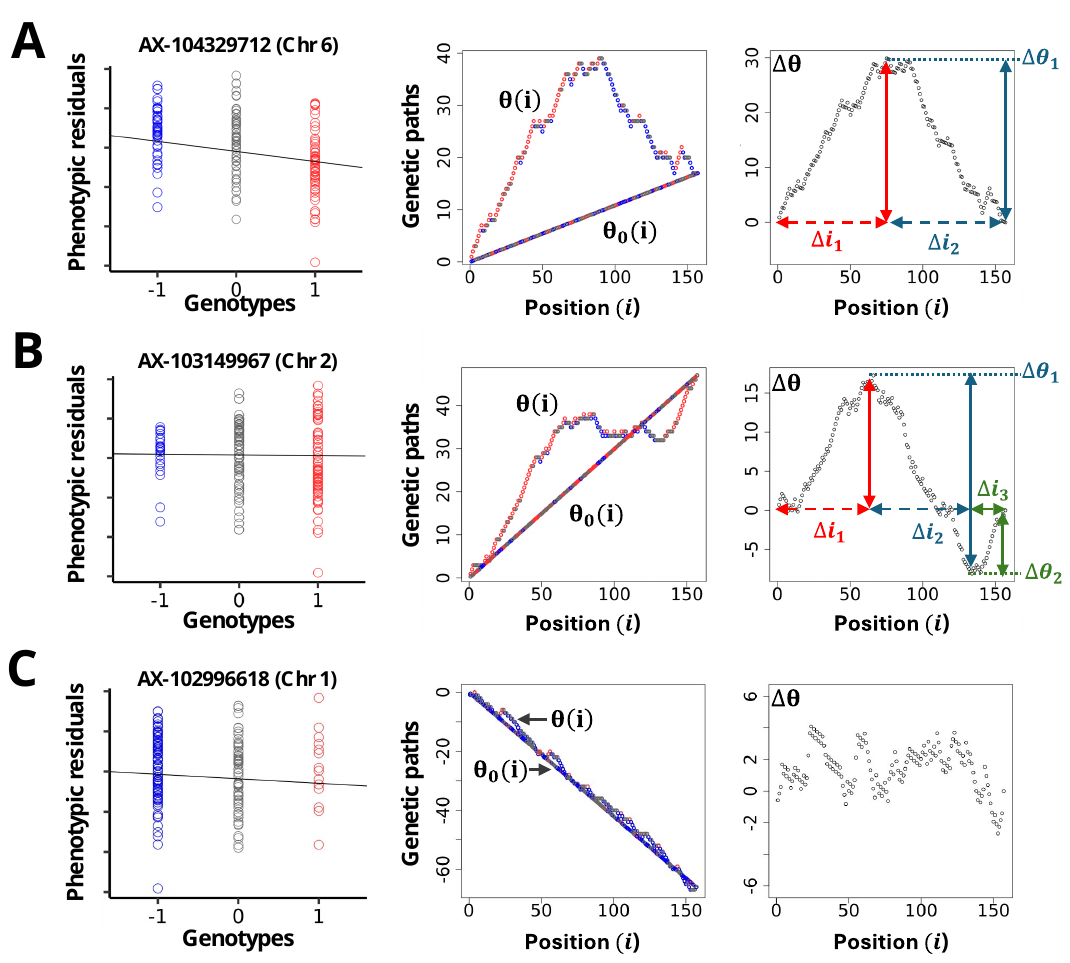

**Figure 2.** Comparison of SNP-Phenotype Associations Using GWAS and GIFT: Three SNPs are represented, illustrating potential genotype-phenotype associations analyzed through GWAS and GIFT. *(A)* SNP associated with the phenotype in both GWAS and GIFT. *(B)* SNP not associated with the phenotype in GWAS but identified as associated in GIFT. *(C)* SNP not associated with the phenotype in either GWAS or GIFT.

To assess whether the organization of microstates is significant, one can calculate the probability of observing extreme path values. For instance, in Fig.2A (right panel), this involves determining the probability that, starting at the origin, $\Delta\theta$ reaches an amplitude ${\Delta\theta}_{1}$ over an interval of positions ${\Delta i}_{1}$, and subsequently returns to zero over another interval ${\Delta i}_{2}$. For the sigmoidal shape of $\Delta\theta$ (Fig.2B, right panel) the probability calculation would involve $\Delta\theta$ starting at the origin, reaching ${\Delta\theta}_{1}$ over ${\Delta i}_{1}$, transitioning from ${\Delta\theta}_{1}$ to ${\Delta\theta}_{2}$ over ${\Delta i}_{2}$, and finally returning to zero over ${\Delta i}_{3}$. Readers interested in the theoretical derivation of p-values for GIFT can refer to Kyratzi et al. (2024) from the list of references given in the main text.

Using this method, the calculated ${-Log}_{10}\left( p_{\mathrm{GIFT}} \right)$ values (level of significances) for $\Delta\theta$ in Fig.2A (right panel) and Fig.2B (right panel) are 22.1 and 24.3, respectively. These results are very similar and demonstrate that GIFT can extract novel information that remains undetectable with GWAS.

Finally, certain SNPs show no significant association with the phenotype using either GWAS or GIFT. This is exemplified in Fig.2C where the SNP yields the p-values of, ${-Log}_{10}\left( p_{\mathrm{GIFT}} \right)=1.406$ and ${-Log}_{10}\left( p_{\mathrm{GWAS}} \right)=0.048$ ($a=-3.00\cdot{10}^{-8} a.u., d=-5.56\cdot{10}^{-6}$ a.u., $r^{2}=0.004$), for GIFT and GWAS, respectively. For GIFT, SNPs unassociated with the phenotype display genetic paths with low amplitude and erratic, disorganized patterns.

By leveraging genetic paths and this approach to calculating p-values for GIFT, Manhattan plots can be generated to compare the effectiveness of GWAS and GIFT in extracting genetic information.
