## Supplemental Data S2 for "Genomic Informational Field Theory (GIFT) to identify genetic associations of a complex trait using a small sample size"

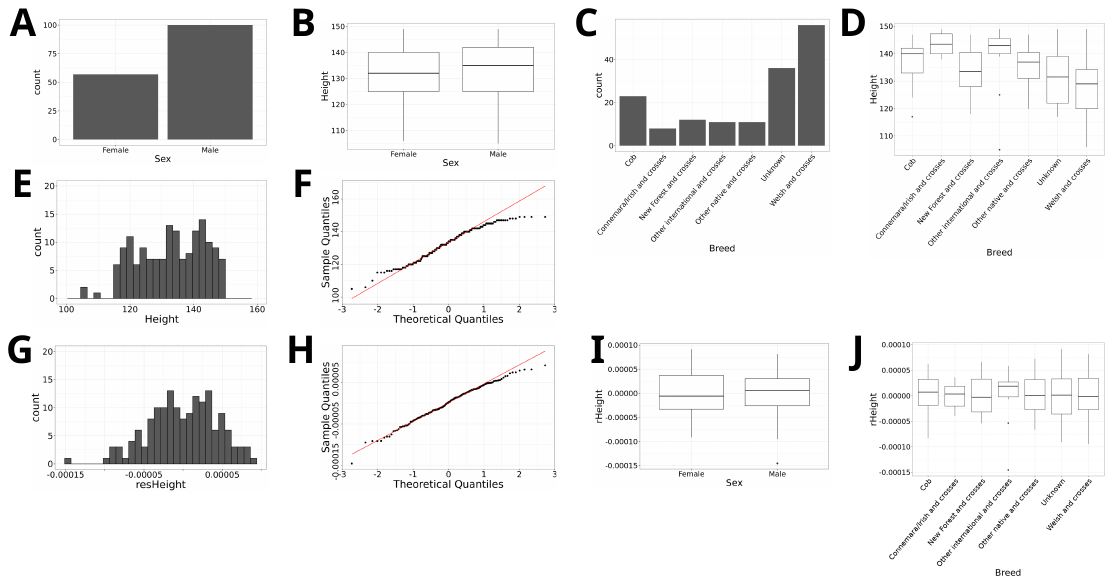

**Extended Data Fig.1: (A)** Counts of female and male in the sample used (n=157). **(B)** Box plots for the distributions of heights at withers measured in cm for female and male. **(C)** Counts of the different breeds involved in the sample used. **(C)** Box plots for the distribution of height at withers per breed measured in cm. **(E)** Counts plot for the heights at withers measured in cm considering the entire sample. **(F)** Quantile-Quantile plot for the height at withers considering raw data (KS-test=0.0510). **(G)** Counts plot of the residual values for the heights at withers (‘resheight’) measured in cm following pre-correction considering the entire sample. **(H)** Quantile-Quantile plot for the residual values for the height withers measured in cm following pre-correction (KS-test=0.8357). **(I)** Box plots of the residual values for the heights at withers (‘resheight’) measured in cm for male and female following pre-correction considering the entire sample (n=157). **(J)** Box plots of the residual values for the heights at withers (‘resheight’) measured in cm across breeds following pre-correction considering the entire sample. (KS-test: Kolmogorov-Simrnov test; cm:centimeter)
